# A Photoactivatable Norepinephrine for Probing Adrenergic Neural Circuits

**DOI:** 10.1101/2023.11.13.566764

**Authors:** Michelle K. Cahill, Yeraldith Rojas Perez, Amara Larpthaveesarp, Roberto Etchenique, Kira E. Poskanzer

**Affiliations:** Neuroscience Graduate Program, University of California, San Francisco, San Francisco, CA 94143, United States; Department of Biochemistry & Biophysics, University of California, San Francisco, San Francisco, CA 94143, United States; Departamento de Química Inorgánica, Analítica y Química Física, INQUIMAE, Facultad de, Ciencias Exactas y Naturales, Universidad de Buenos Aires, CONICET, Ciudad Universitaria Pabellón 2, AR1428EHA Buenos Aires, Argentina; Department of Pediatrics, University of California, San Francisco, San Francisco, CA 94143, United States; Kavli Institute for Fundamental Neuroscience, San Francisco, CA 94143, United States

**Keywords:** Ruthenium complex, norepinephrine, electrophysiology, neuromodulator, phototrigger, uncaging

## Abstract

Norepinephrine (NE) is a critical neuromodulator that mediates a wide range of behavior and neurophysiology, including attention, arousal, plasticity, and memory consolidation. A major source of NE is the brainstem nucleus the locus coeruleus (LC), which sends widespread projections throughout the central nervous system (CNS). Efforts to dissect this complex noradrenergic circuitry have driven the development of many tools that detect endogenous NE or modulate widespread NE release via LC activation and inhibition. While these tools have enabled research that elucidates physiological roles of NE, additional tools to probe these circuits with a higher degree of spatial precision could enable a finer delineation of function. Here, we describe the synthesis and chemical properties of a photo-activatable NE, [Ru(bpy)_2_(PMe_3_)(NE)]PF_6_ (RuBi-NE). We validate the one-photon (1P) release of NE using whole-cell patch clamp electrophysiology in acute mouse brain slices containing the LC. We show that a 10 ms pulse of blue light, in the presence of RuBi-NE, briefly modulates the firing rate of LC neurons via α-2 adrenergic receptors. The development of a photo-activatable NE that can be released with light in the visible spectrum provides a new tool for fine-grained mapping of complex noradrenergic circuits, as well as the ability to probe how NE acts on non-neuronal cells in the CNS.

## Introduction

The neuromodulator norepinephrine (NE) plays fundamental roles in the nervous system by mediating attention^1–3^, plasticity^4,5^, and memory consolidation^6^. Further, an imbalance in NE levels has been implicated in both attention-deficit/hyperactivity disorder^7,8^ and depression^9–11^. NE performs this wide range of neuromodulatory functions by acting on a number of receptors across diverse cell types in the central nervous system, including neurons, astrocytes, and microglia^12–20^. Given both the fundamental roles of NE in the central nervous system (CNS) and the diversity of its cellular targets, tools to dissect this complex neuromodulatory system are critical to understand its many functions. A recent push to dissect noradrenergic circuit logic has led to the development of NE-based tools both to detect endogenous NE activity and to experimentally stimulate or inhibit NE release. Development of fluorescent sensors for NE, such as GRAB_NE_ ^21^ and a carbon nanotube-based NE nanosensor^22^, allow for *in vivo* detection of endogenous NE release. In an effort to manipulate noradrenergic circuits, a number of groups have optogenetically activated or inhibited the locus coeruleus (LC) to either stimulate^23–25^ or inhibit^23,26^ NE release throughout the CNS. While optogenetic activation of the LC provides a physiologically relevant spatial pattern of release, this spatial pattern is inherently widespread, targeting many regions of the CNS^27–29^. Due to their highly collateralized nature, even targeted stimulation of a single axon or neuron may result in NE release across a relatively broad field^29,30^. Additionally, it has been shown that LC neurons have the capacity to co-release dopamine along with NE^31^. Thus, to provide a tool to probe noradrenergic circuits with a higher degree of spatial precision and to isolate the specific action of NE on cells and circuits, we have developed a photoactivatable NE.

Photoactivatable compounds have proven to be powerful tools for mapping functional connectivity of GABAergic and glutamatergic circuits^32–34^, receptor localization^35^, and locations of synaptic inputs onto subcellular compartments of neurons^36^. A photoactivatable compound is a molecule “caged within” a protecting group that is released upon absorption of light. This reaction occurs quickly, and the amount of molecule released can be scaled by altering light intensity and beam size. Thus, photoactivatable compounds make it possible to release molecules in a temporally and spatially precise manner, without mechanically disturbing the preparation. A photoactivatable NE has previously been developed^37^. However, it is activated by light in the UV spectrum, which can be damaging to, and have difficulty penetrating into, biological tissue. Thus, a new photoactivatable NE, activated by light in the visible spectrum, would greatly improve our ability to probe noradrenergic circuits in the same way that photoactivatable glutamate and GABA have enabled a detailed mapping of their respective circuits and receptors.

Ruthenium-bipyridine caged compounds were first presented two decades ago^38^ as a new way to extend the activation spectrum to longer wavelengths. These cages are comprised of a ruthenium core coordinated to one or two bipyridines or analogue molecules, and one or two monodentate ligands. These complexes show a strong absorption in the visible region (usually blue-to-green), corresponding to a ^1^MLCT charge transfer band^39^. Upon irradiation of such band, in a fast sequence that occurs in tens of nanoseconds, a dissociative d-d state is populated, which leads to the release of one of the monodentate ligands, usually the molecule of interest. The remaining monodentate ligand can be used to tune the chemical and photochemical properties of the complex.

Here, we describe the synthesis and chemical properties of RuBi-NE, which releases NE upon absorption of visible light. Additionally, we demonstrate its use in a biological system by modulating the firing rate of locus coeruleus (LC) neurons, neurons known to be autoregulated by NE^40,41^. In the future, RuBi-NE allows for the stimulation of cells and circuits with NE in a spatially and temporally precise manner. Targeted release of NE will allow for a fine-grained mapping of the functional connectivity of noradrenergic circuits, as well as an ability to probe how NE acts on non-neuronal cells in the CNS. The ability to dissect noradrenergic circuits with this level of precision will help to move the field toward a deeper understanding of how NE performs its wide range of neuromodulatory functions.

## Results

Although the synthesis of RuBi caged compounds is straightforward^42^, special care must be taken to obtain RuBi-NE, due to the tendency of free NE phenol groups to undergo oxidation in the basic media needed for coordination. Thus, the synthesis must be performed in deep anaerobic conditions using Schlenk procedures (see Methods). Once obtained, RuBi-NE is a stable solid that can be stored at room temperature, although a -15°C freezer is preferable for long-term storage. RuBi-NE is a +2 charged cation at low pH, although in physiological conditions its phenol groups can be partially deprotonated.

To confirm the biocompatibility of the RuBi-NE complex and evaluate its performance as a NE phototrigger, a series of electrophysiology experiments in acute brain slices containing LC were performed. First, we identified LC neurons for recording by crossing a mouse line that expresses Cre in LC neurons (*TH-Cre*) with a Cre-inducible fluorescent reporter line (*Ai14*) to drive expression of the red-shifted fluorescent molecule TdTomato specifically in LC neurons (Fig 1a). These fluorescently labeled neurons exhibited the stereotypical large morphology of LC neuronal somata. Next, we targeted these cells for whole-cell patch-clamp recordings, including Alexa dye in the pipette solution to confirm that the correct cell was patch-clamped (Fig 1a). Patched cells were held in the current clamp configuration, and we observed that these neurons spontaneously fired action potentials (APs), as has been described previously^43–46^. We took advantage of the fact that LC neurons themselves express α_2_ adrenergic receptors (ARs), and NE activation of these receptors acts to inhibits AP firing of these cells^41^. We confirmed this cellular behavior by bathing the slice with 30μM NE, and observed a marked decrease in AP firing, as expected (Fig 1b). We thus reasoned that any increase in NE at the patched cell would decrease AP firing.

**Figure 1.**
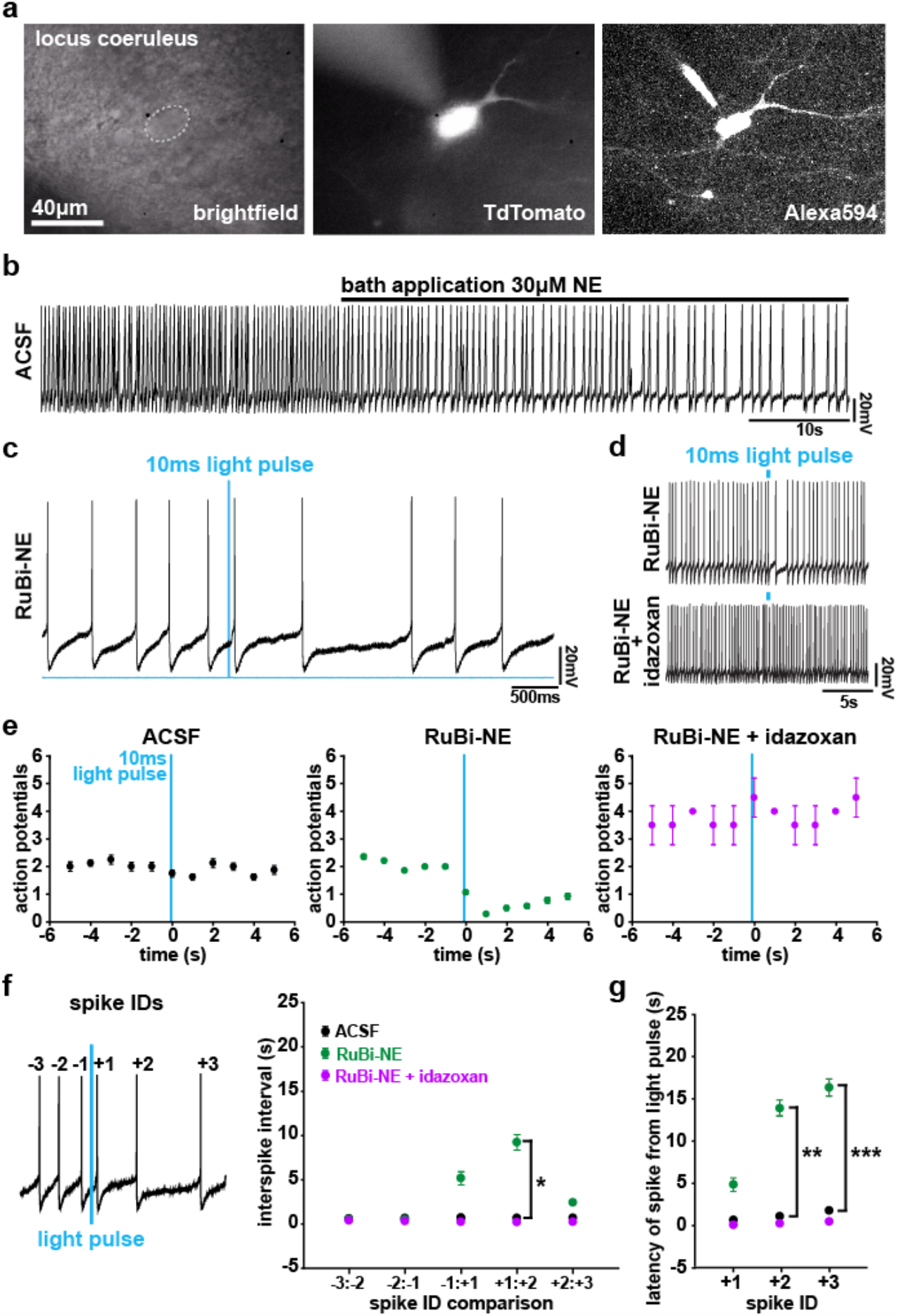
Photorelease of RuBi-NE specifically increases local [NE]. (a) Representative image of a LC neuron identified for recording. Left: DODT contrast image of an LC neuron before recording; dotted line marks soma. Center: 1P image of tdTomato^+^ LC neuron before whole-cell patch-clamp configuration. Right: 2P image of the same LC neuron in whole-cell patch-clamp configuration, dialyzed with Alexa 594. Patch pipette shown on the upper left border of the cell. (b) Representative trace of spontaneous AP firing pattern of LC neuron in current clamp configuration. Bath application of NE (30 μM) occurs 30 s into the trace (black bar). (c) Representative activity of an LC neuron before and after uncaging RuBi-NE. Blue line marks 10 ms blue-light uncaging pulse. (d) Zoomed out representative activity before and after RuBi-NE uncaging before (top) and during (bottom) blockade of α_2_ ARs with idazoxan (2 μM). Blue rectangles mark 10 ms blue-light pulse. (e) Average APs/s for an LC neuron in ACSF (n = 8 pulses in 2 cells, left), in 300 μM RuBi-NE (n = 14 pulses in 4 cells, center), and in 300 μM RuBi-NE + 2 μM idazoxan (n = 2 pulses in 1 cell, right). *Time 0* denotes the second in which the light pulse occurred. (f) Left: Example trace showing spike IDs relative to the light pulse. Right: Average ISI calculated for LC neurons in ACSF (n = 8 pulses in 2 cells), RuBi-NE (n = 14 pulses in 4 cells) and RuBi-NE + idazoxan (n = 2 pulses in 1 cell). *-1:+1* is the ISI during which uncaging occurs. Two-way ANOVA grouped by condition and ISI number followed by Bonferroni test to determine significant pairwise comparisons, * p < 0.05. (g) Average absolute time of the first three spikes following blue-light pulse in ACSF, RuBi-NE, and RuBi-NE + idazoxan. Two-way ANOVA grouped by condition and spike number followed by Bonferroni test to determine significant pairwise comparisons, ** p < 0.01 and *** p < 0.001.

To test whether photoactivation of RuBi-NE causes increased local NE, we added RuBi-NE to the bath in standard artificial cerebraspinal fluid (ACSF), without any additional NE. We first confirmed that RuBi-NE in the bath solution did not change cellular activity of the patched neuron, with AP firing rate unchanged between the two conditions (ACSF: 1.94 ± 0.31 spikes/s and RuBi-NE: 2.09 ± 0.10 spikes/s, p = 0.60, unpaired t-test). However, upon one-photon (1P) activation (10 ms) using 446–486 nm blue light, we observed a consistent decrease in AP firing rate (Fig 1c, d [top], e [middle]), indicating that NE was indeed photoreleased. The blue light itself did not have any effect on AP firing, since the same effect was not observed when RuBi-NE was not added to the bath (Fig 1e, left). Next, to confirm that the change in AP firing was due to release of NE and activation of ARs, we carried out the same 1P-photoactivation but added the α_2_ AR antagonist (idazoxan, 2 μM) to the bath along with RuBi-NE. Upon blue-light activation in these experiments, we no longer observed a change in AP firing (Fig 1d [bottom], e [right]), although idazoxan itself increased firing rate, as previously^43,47^. These data indicate that RuBi-NE photoactivation specifically increases local NE concentration.

To demonstrate the temporal dynamics of the NE photoactivation on LC neurons, we also calculated the interspike interval (ISI) between the APs occurring before and after the 10 ms blue-light pulse. When quantifying the ISI, we detected no significant difference among conditions before the blue light photoactivation (RuBi-NE: 0.5151 ± 0.0082s, 0.4984 ± 0.007s, ACSF: 0.5992 ± 0.0578s, 0.6453 ± 0.0545s and RuBi-NE + idazoxan: 0.3794 ± 0.1227s, 0.3071 ± 0.0473s, for each of the two ISIs, respectively, immediately preceding the light pulse), but a significant delay between spikes after the pulse with RuBi-NE in the bath compared to the ACSF condition (RuBi-NE: 5.1641 ± 0.7551s, 9.2122 ± 0.8788s, 2.45165 ± 0.1469s, ACSF: 0.6735 ± 0.0454s, 0.6549 ± 0.0424s, 0.6708 ± 0.0516s and RuBi-NE + idazoxan: 0.2201 ± 0.0109s, 0.1901 ± 0.051s, 0.2252 ± 0.0226s, for each of the three ISIs respectively, including and immediately following the light pulse. Two-way ANOVA followed by Bonferroni test determine significant pairwise comparisons between conditions and ISIs; *p* = 0.0492 between RuBi-NE and ACSF between spikes 1:2 following the light pulse) (Fig 1f, green). These ISI delays were not observed when the experiment was carried out in ACSF only (Fig 1f, black), or when idazoxan was applied to the bath with RuBi-NE (Fig 1f, pink). Similarly, when we quantify the absolute time of each AP from the light pulse, we find that a 10 ms photo-release of RuBi-NE significantly delays the firing of subsequent APs, such that the second and third AP after the light pulse occurs 14.0546 ± 0.9813s and 16.47 ± 1.0317s after the light stimulation (Fig 1g, green), while the control APs after the pulse occurred at 1.05 ± 0.0603s and 1.73 ± 0.1042s, and 0.3068 ± 0.1407s and 0.53 ± 0.1633s (Fig 1g, black and pink, respectively, two-way ANOVA followed by Bonferroni test determine significant pairwise comparisons between conditions and APs; *p* = 0.0044 and *p* = 0.00040 between RuBi-NE and ACSF at spike +2 and +3, respectively). In all, these data indicate that RuBi-NE acts as a robust, specific phototrigger for NE.

## Discussion

Here, we introduce the development of [Ru(bpy)_2_(PMe_3_)(Norepinephrine)]^n+^ (RuBi-NE), a photoactivatable NE releasable with light in the visible spectrum. We validate the release of NE with one-photon uncaging in acute mouse brain slices containing the LC. We find that uncaging RuBi-NE, with a brief pulse of blue light, temporarily slows the rate of LC action potentials via α-2 adrenergic receptors. This new tool will allow for the temporally and spatially precise release of NE without mechanical disturbance of tissue, which itself can modulate cellular activity^48^.

Our validation of 1P NE uncaging releases NE over a large field-of-view, the size of the uncaging light beam. This widespread release of NE is physiologically relevant given the extensive projections of LC neurons throughout the brain, which release NE via volume transmission^49,50^. While this spatially diffuse uncaging allows for investigation of the specific action of NE isolated from any potential co-release from synaptic terminals, validation of the two-photon compatibility of RuBi-NE, as demonstrated for other RuBi subtypes^51,52^, will be needed to map noradrenergic circuitry with subcellular spatial precision. Synthesis and validation of this photoactivatable compound adds NE to the suite of neurotransmitters and neuromodulators that can now be caged within a ruthenium-bipyridine backbone, expanding the toolkit available to map functional circuitry and receptor localization of neuronal and non-neuronal cells in the central nervous system.

## Methods

### RuBi-NE synthesis

All chemicals were purchased from Sigma-Aldrich and used as received without further purification. Ru(bpy)_2_Cl_2_ was synthesized according to literature^53^. From this, [Ru(bpy)_2_(PMe_3_)(Cl)]PF_6_ was obtained as previously described^54^. For synthesis of [Ru(bpy)_2_(PMe_3_)(Norepinephrine)]^2+^, 300 mg of [Ru(bpy)_2_(PMe_3_)(Cl)]PF_6_ was dissolved in 3 ml acetone and 15 ml water was added. The mixture was stirred overnight with mesylate-loaded Dowex-22 ion exchange resin. After evaporation of acetone, the resultant solution of [Ru(bpy)_2_(PMe_3_)(H_2_O)]Mes_2_ was separated from the resin. From this point all procedures were performed under N2, with the solution previously degassed with Ar or N_2_. 450 mg of norepinephrine hydrochloride was introduced into a Schlenk flask, and the pH was raised to 9–10 by adding 2 mL of 1 M NaOH. When the formation of the NE complex was confirmed by UV-visible spectroscopy, the mixture was cooled in an ice bath, and 1 M acetic acid was added to lower the pH and prevent possible oxidation of NE by air during the subsequent procedures. The solution was filtered, and after the addition of 2 mL of 0.5 M KPF_6_, the complex [Ru(bpy)_2_(PMe_3_)(NE)](PF_6_)_2_ precipitated as an orange powder. For a more soluble preparation, a further ion-exchange with mesylate-loaded Dowex 22 resin can be made.

### Slicing and electrophysiology

Horizontal brain slices (350 μm) at the level of the pons in the brainstem containing the LC were prepared from P28–P34 TH-Cre^+^; Ai14^+^ and TH-Cre^+^; Ai9^+/-^ female mice using a vibratome (VT1200; Leica). Slicing solution was chilled at -20°C and contained (in mM): 222 sucrose, 27 NaHCO_3_, 2.6 KCl, 2 MgSO_4_, 2 CaCl_2,_ 1.5 NaH_2_PO_4_, bubbled with 95% O_2_/ 5% CO_2_. Slices were incubated in artificial cerebrospinal fluid (ACSF) at 35°C for 30 min and then held at room temperature until the time of recording. Recording ACSF contained (in mM): 123 NaCl, 26 NaHCO_3_, 10 dextrose, 3 KCl, 2 MgSO_4_, 2 CaCl_2_, 1 NaH_2_PO_4_, bubbled with 95% O_2_/5% CO_2._.

Neurons were visualized using DODT contrast microscopy on an upright microscope (Bruker) and fluorescently labeled tdTomato-positive LC neurons were identified using an EXFO X-Cite 120 epifluorescent lamp (Excelitas) coupled to a red filter cube (TRITC-B-000, Semrock). Neurons were visualized using a NIR Apo 40X/0.80W objective (Nikon) and an infrared CCD real-time camera (IR-2000, DAGE-MTI). Recordings were made using a Multiclamp 700B (Molecular Devices) amplifier and acquired at a rate of 10 kHz with Prairie View 5.4 software. Patch pipettes (6–7 MΩ tip resistance, borosilicate glass, outer diameter 1.5 mm, inner diameter 0.86 mm, Sutter Instruments) were filled with the following internal solution (in mM): 135 K-methylsulfate, 10 KCl, 10 HEPES, 5 NaCl, 0.025 Alexa 594, pH 7.3 (adjusted with KOH). Series resistance ranged between 15–30 MΩ, and pipette capacitance was compensated. Neurons were recorded in current clamp configuration. With no injected current, neurons had a resting membrane potential (V_m_)∼ -50mV.

### RuBi-NE uncaging

Once the neuron was in whole-cell patch clamp configuration, all lights were turned off and a stock solution of 10mg/mL RuBi-NE in ACSF was added to the bath solution for a final RuBi-NE concentration of 300 μM. While recording, RuBi-NE was photolyzed by 1P excitation with a single 10 ms pulse of blue light (446–486 nm) using an EXFO X-Cite 120 epifluorescent lamp (Excelitas) coupled with a 466/40 nm single-band bandpass filter (BrightLine, Semrock) at an approximate power of 14.3mW. Idazoxan hydrochloride (Sigma Aldrich, 2 μM) was added to the ACSF bath solution to confirm that RuBi-NE was acting on the neurons via α_2_ adrenergic receptors.

### Statistics

Averages are reported as mean ± SEM. Average firing rates before the light pulse were calculated by taking the mean firing rate across pulses. An unpaired t-test was used to calculate the p-value to compare the average firing rate before the light pulse between the ACSF and RuBi-NE conditions. The remaining p-values were calculated using two-way ANOVAs followed by Bonferroni tests determine significant pairwise comparisons between conditions and time points.

## Acknowledgements

K.E.P.’s work was supported by NIH R01NS099254, NSF 1604544, and the Brain Research Foundation Frank/Fay Seed Grant. M.K.C. was supported by funding from the NSF Graduate Research Fellowship 1650113. R.E. is a member of CONICET. R.E. thanks ANPCyT and UBACyT for funding. We thank Jennifer Thompson for administrative support. This material is based upon work supported by the National Science Foundation Graduate Research Fellowship Program under Grant No. 1650113. Any opinions, findings, and conclusions or recommendations expressed in this material are those of the author(s) and do not necessarily reflect the views of the National Science Foundation.

